# A cell based, high throughput assay for quantitative analysis of Hedgehog pathway activation using a Smoothened phosphorylation sensor

**DOI:** 10.1101/110056

**Authors:** Eugene A. Albert, Christian Bökel

## Abstract

The Hedgehog (Hh) signalling cascade is conserved across evolution and plays an important role in development and disease. In the absence of Hh, activity of the key signal transducer Smoothened (Smo) is downregulated by the Hh receptor Patched (Ptc). However, the mechanisms underlying this inhibition, and especially its release upon ligand stimulation, are still poorly understood, in part because tools for directly following Smo activation at the subcellular level were long lacking. Here we present a high throughput, cell culture assay based on a fluorescent sensor for *Drosophila* Smo phosphorlyation. Using this approach we could first demonstrate that the graded response to increasing Hh levels observed at the population level can be traced back to threshold responses of individual cells exhibiting differential Hh sensitivity. Second, we screened a small molecule inhibitor library for regulators of Smo phosphorylation. We observed increased Smo sensor fluorescence with compounds aimed at two major target groups, the MAPK signalling cascade and polo and aurora kinases. Biochemical validation confirmed the screen results for selected inhibitors (dobrafenib, tak-733, volasertib) and revealed differences in the mode of Smo activation, demonstrating that the assay is in principle suitable for dissecting the cell biological basis of Hh pathway activation.

## Introduction

Hedgehog (Hh) signalling plays an important role in development and disease, and is highly conserved across different branches of the evolutionary tree. A unique feature of the Hh signalling cascade is the sequential use of two receptor-like proteins, the actual Hh binding receptor Patched (Ptc) and the downstream, GPCR-like signal transducer Smoothened (Smo). In the absence of Hh, Ptc suppresses the activity of Smo, retaining it in an endosomal compartment. Upon Hh binding to Ptc, this suppression is released, leading to Smo translocation to plasma membrane and activation of the downstream signalling cascade. However, while the downstream events in Hh signal transduction are reasonably well understoood, the mechanisms underlying the Ptc-mediated suppression of Smo activity, and the upstream events leading to Smo activation during pathway activation, remain to be fully elucidated despite almost 30 years of research into the Hh pathway ^1^.

Since Ptc is structurally a member of the RND family of small molecule transporters ^2^, it has been suggested to act as a transporter for small molecules that influence Smo activity. Extensive efforts have therefore been focused on identifying endogenous Smo ligands that may be trafficked by Ptc. In vertebrates, sterols were implicated through a line of evidence that started with the importance of cholesterol metabolism for signal transduction ^3, 4^, continued with a detailed biochemical model ^5^ and just recently was extended to structural biology ^6, 7^. However, a role of sterols as the main link between Ptc and Smo is still under debate, especially in *Drosophila*, where the pathway was initially discovered. Instead, endocannabinoids were proposed as alternative, potential Smo ligands that may act as suppressors of Smo activity both flies and vertebrates ^8^ and may thus coordinate Hh signalling at the cellular and organismic level. However, it is not clear whether these endocannabinoids are the primary targets of Ptc regulation. Phospholipids are a third class of small molecules suggested to be affecting Smo downstream of Ptc activity. In *Drosophila*, increased PI4P levels were shown to promote Hh signalling ^9^. More recent data provided evidence for the direct regulation of phospholipids by Hh and binding of PI4P to Smo ^10^. Nevertheless, none of these molecules are widely accepted to be the major, Ptc dependent Smo regulators.

A similar research effort was focused on describing the molecular events occurring at the level of Smo during pathways activation. Most prominently, phosphorylation of *Drosophila* Smo by PKA primes it for further phosphorylation by the CK and GPRK kinases ^11, 12^. Phosphorylation protects Smo from ubiquitination by interfering with ubiquitin ligases and through the recruitment of deubiquitinating enzymes ^13, 14^. Since ubiquitination promotes internalization of Smo, phosphorylation stabilizes active Smo at the plasma membrane ^13, 14^, resembling the effect of sumoylation ^15^. Since Smo has to be present at the plasma membrane in order to activate downstream pathway components, endocytosis also plays an important role in Hh pathway regulation. Indeed, trapping Smo on the plasma membrane is sufficient to promote Smo phosphorylation, potentially placing Smo localization upstream of Smo phosphorylation ^16^.

However, despite all these individual advances in the field, we are still lacking a comprehensive picture of the early events in Hh pathway activation. Unfortunately, screening specifically for upstream mechanisms affecting Smo activation has, to date, been difficult. Several general screens using transcriptional readouts have identified additional components of the Hh cascade, thus providing valuable insight in our understanding of the system ^17–21^. Nevertheless, this strategy also has limitations. Most prominently it responds to the final outcome of pathway activation. It is therefore likely to miss events that partially perturb Smo activation but whose effect on gene expression may be buffered or masked by downstream components of the cascade, e.g. through signal amplification and feedback mechanisms. A system that would allow us to directly follow Smo activation, uncoupling it from both external influences and internal feedback processes, could potentially help to shed light on this question.

We have previously described a fluorescence based sensor (SmoIP) that can visualize endogenous or experimental phosphorylation of *Drosophila* Smo in transgenic flies ^16^ by detecting the associated disruption of an off-state specific intramolecular loop in the Smo cytoplasmic tail ^22^. For this, the circularly permutated GFP (cpGFP) core of the Inverse Pericam Ca^2+^ sensor ^23^ was inserted into the C-terminal Smo cytoplasmic tail such that the formation of the intracellular loop forces the cpGFP into an nonfluorescent state, while the release of the loop by phosphorylation lets the cpGFP core relax into a fluorescent conformation (Fig. 1a). Using this sensor in a screening setup could potentially overcome the limitations of the transcriptional readouts, and would allow to systematically investigate pathway activation at the receptor level.

**Figure 1.**
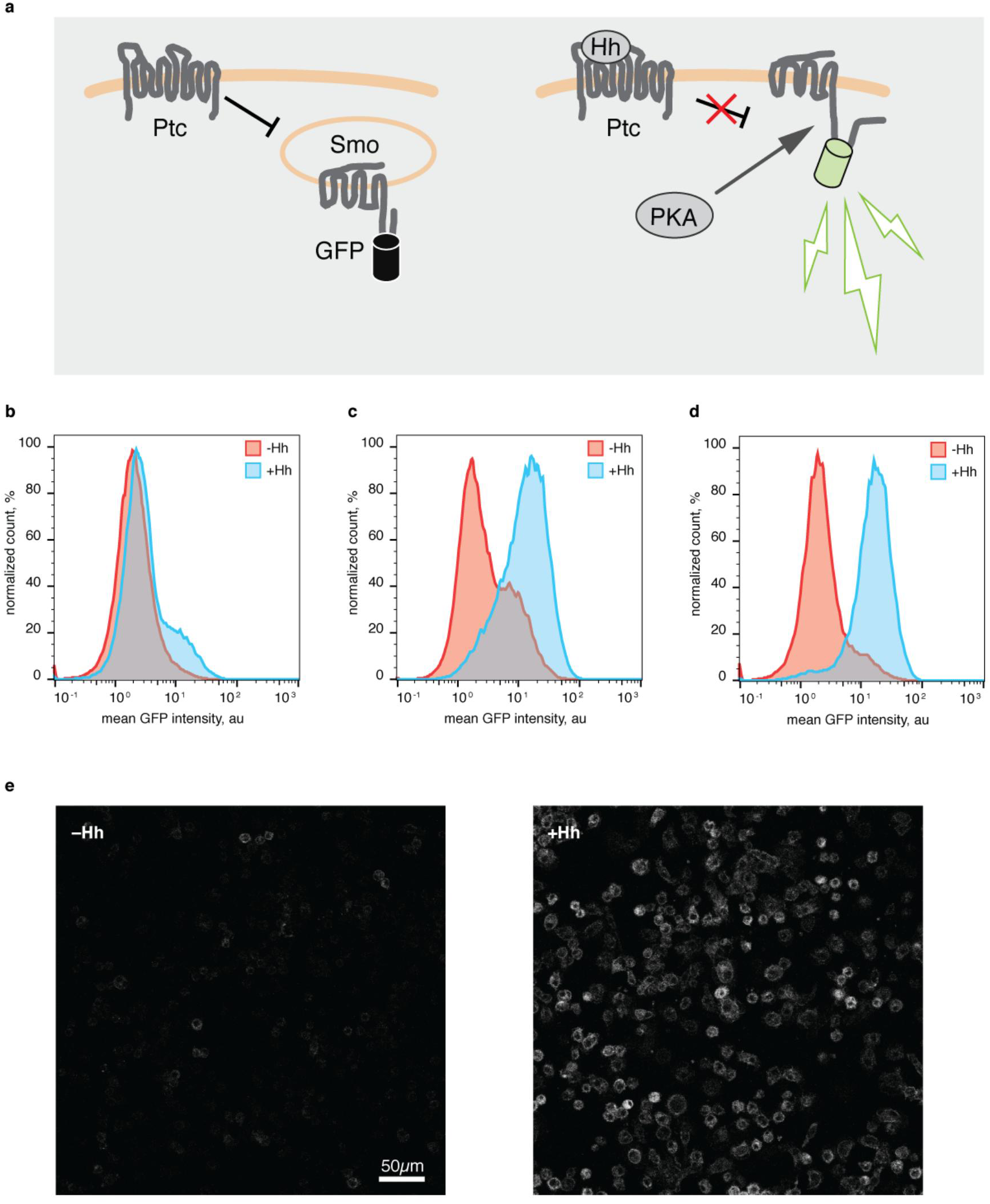
Direct detection of Smo phosphorylation using a SmoIP transgenic S2 cell line. **(A)** SmoIP detection principle. In the inactive state, an intramolecular loopin the Smo cytoplasmic tail forces an inserted cpGFP cassette from the Inverse Pericam (IP) Ca^2+^ sensor into a nonfluorescent conformation. During pathway activation, dissolution of this loop by Smo phosphorylation allows the IP cassette to relax into a fluorescent state. (B-D) FACS analysis of UAS-SmoIP transgenic cell lines following 24h stimulation with Hh conditioned medium. Cells were screened as a polyclonal line following 3 weeks selection (B), following iterative sorting of strongly responding cells (C), and as a single cell derived, clonal line (cl14) (D). (E) Confocal live-cell images of cl14 response to Hh. Scale bar 50*µ*m.

Here, we report the development of a cell-based assay based on the SmoIP sensor for the direct, quantitative, high throughput assessment of the Smo phosphorylation state. Using this system we could first demonstrate that individual cells within a largely homogenous, clonal population activate Smo at cell specific threshold concentrations of Hh. Second, we performed a proof of principle experiment, testing a library of small molecule inhibitors with defined molecular targets for their effects on Smo activation. We thereby identified components of the MAPK pathway as well as the polo and aurora kinases involved in cell cycle regualtion as two major clusters of target proteins, whose inhibition affects Smo phosphorylation through distinct molecular mechanisms. These observations thus provide proof of principle for the suitability of our assay for high throughput screening experiments, and provide novel insights in the logic of Smo activation.

## Methods

### Plasmids

The P-element vectors pUAST or pCasPer using the appropriate inserts were used for the generation of cell culture plasmids. To generate the UAS: smoIP plasmid, the white^+^ transgenesis marker was of pUAST-SmoIP was replaced with act: Hygro and tubulin: Gal4VP16 cassettes. For tubulin: smoIP, the act: DHFR selection cassette was cloned into the backbone of pCasPer4-tubuin: SmoIP in place of the white^+^ marker. For mtn: HhN, an act: Hygro selection cassette was cloned into the backbone of a pCasPer4 plasmid carrying a mtn: HhN insert. Detailed plasimds maps are available upon request.

### Cell culture

S2 cells were cultivated at room temperature in Schneider´s Drosophila Medium (Pan Biotech) supplemented with 10% FBS (ThermoFisher) and pen/strep antibiotics in T25 flasks (TPP). Cell transfection was done using a standard calcium phosphate protocol ^24^. For P-element transformation, target plasmid was co-transfected with ∆2−3 helper plasmid ^25^ at a 10:2 ratio. Stable cell lines were selected on either 300*µ*g/ml hygromycinB (Sigma-Aldrich) or methotrexate 4*10^−7^ M (Sigma-Aldrich) for 3 weeks ^26^. Clonal selection was done according to protocol ^27^. Briefly, S2 feeder cells were irradiated with 23.3 kR. Target cells were diluted to a concentration of 50 cells/ml, mixed with feeder cells (10*10^6^) in 8ml of full growth medium. 2ml of 1.5% agarose (filter sterilized) were added to the cells and mixed by gentle shaking. Individual clonal colonies were excised in 2−3 weeks after reaching 2−3mm in diameter, the surrounding agar was mechanical softened and cells were placed in 96 well plates (ThermoFisher). Expression from the mtn promoter was induced using 1mM of CuSO_4_. Hh condition medium was collected from cells after 7d of growth, filter-sterilized and stored at 4°C for up to 5 months. Mock medium was produced in the same fashion by wild type S2 cells. For all experiments except screens and western blots cells were stimulated in 96 well plates at a density of 10^5^ cells / well with 20*µ*L of conditioned medium. Fluorescence was measured after 24h not specified otherwise. For ds RNA experiments cells were seeded on 96 well paltes in CCM3 medium (ThermoFisher) at a density of 30*10^3^ cells in 50*µ*L containing 2*µ*g dsRNA (Sheffield iRNA screening facility) per well. Cells were analyzed 4 days later.

Inhibitors were used in the following concentrations: OA, 5 nM; IBMX, 24*µ*g/ml, fsk, 80*µ*M; h89, 30*µ*M, dbn, tak-733 and vlt, 15*µ*M if not stated otherwise.

Cl8 cells were cultivated under standard conditions in M3 medium (Sigma-Aldrich, USA), supplemented with 2.5% FBS (ThermoFisher, USA), and 2.5% fly extract (prepared according to the DGRC protocol), insulin at 5*µ*g/ml and pen/strep solution.

### FACS

Cells were analyzed either by MACSQuant (Miltenyi biotec) or by FacsCanto II analyzers (BD). Fluorescence was acquired directly in the growth medium after cells detachment by pipetting. Cell sorting was done on a facsAria II sorter (BD).

### Screening

Cl14 cells were dispensed to 384 well V-bottom plates (Eppendorf) containing a pre-aliquoted library of small molecule inhibitors (Selleckchem, Houston, USA, library L1100) at concentrations of either 2*µ*M or 15*µ*M, in 25*µ*l of medium at a density of 25*10^3^ cells per well. After 30min of incubation cells were stimulated with 5*µ*L of Hh conditioned or mock medium. After 24h, 4000 cells per well were analyzed using a BD cantoII FACSanalyzer.

### Western blots

3*10^6^ S2 cells were seeded on 6 well plates in 1,5 ml volume and stimulated with 500*µ*L of either Hedgehog or mock conditioned medium for 24h hours. Cells were lysed in lysis buffer (25 mM Tris pH 7.2, 150 mM NaCl, 5 mM MgCl2, 0.2% NP-40, 1 mM DTT, 5% glycerol) on ice in presence of protease (Roche) and phosphatase (Sigma-Aldrich) inhibitor cocktails for 30min. Lysates were boiled for 20sec with loading buffer, cooled on ice an run on bis-tris gradient gels (ThermoFisher). Ptc (Apa 1, 1:250), smo (20C6, 1:100), and tub (12G10, 1:250) antibodies were obtained from DSHB, beta-act (1:5000) from Abcam, GFP antibody was obtained from Clontech. The HhN antibody (1:500) was a gift of Suzanne Eaton (Dresden). Primary antibody incubation was performed at 4°C overnight, secondary antibody incubation for 1h at RT.

### Plasma membrane smo staining

After stimulation for 24h with Hh conditioned medium 2*10^6^ cl14 cells were harvested by pipetting and centrifugation, re-suspended in 150uL FACS buffer (0.1% NaN_3_, 0.5% BSA in PEM) containing Smo ab (1:300) and stained for 1h 30min at RT on shaker. Then cells were centrifuged, washed once with 2ml of ice-cold FACS buffer for 10 min, centrifuged and stained with AlexaFluro 647 tagged secondary antiserum (1:500) for 30 min at RT. Cells were then centrifuged, washed, and centrifuged again, re-suspended in 200*µ*L of FACS ice-cold buffer and analyzed by FACS.

### Real time qPCR

3*10^6^ of S2 cells were treated with inhibitors in 2ml medium in 35mm petri dishes for 24h before RNA extraction. RNA was extracted using Trizol (ThermoFisher). cDNA was synthesized using a first strand cDNA synthesis kit (ThermoFisher) and random primers from 3*µ*g of RNA. qPCR was performed with maxima SYBR green qPCR master mix (ThermoFisher) on a lightCycler 480 (Roche) using the following primer pairs:

ptc_F: AGTCCACGAACAATCCGCA

ptc_R: TGGGTCGTCTGAATGAGCAG

gapdh2_ F: GAGTTTTCGCCCATAGAAAGC

gapdh2_R: CGATGCGACCAAATCCATTG ^28^

Ptc expression was calculated using the ddCt method relative to gapdh2 expression.

### Microscopy

Cells were live-imaged 24h after stimulation with Hh condition medium. 2-3h prior to imaging cells were lifted by pipeting and placed in Lab-Tek chambers (ThermoFisher). Images were taken using Zeiss LSM780 confocal microscope and analyzed by ImageJ.

### Data analysis

FACS data was analysed by FlowJo (FlowJo Software, USA). Gating for FSC and SSC axis in each experiment was done according to the control cells. For the screen results, only samples with more than 50% of cells in the chosen gate were taken in to account. Median GFP intensity was calculated for the gated population in each sample. Treatment effects were measured as a difference between signal intensity of inhibitor and a non-stimulated control normalized to the difference between stimulated and non-stimulated controls, hereafter referred to as “normalized response”. Analysis and visualization of the screen results was done with the help of the Pandas Python package. All charts represent mean ± sd values with sample sizes indicated in figure legend. Significance levels were calculated using the Mann-Whitney U test implemented in the ScyPy Python package.

## Results

*The cell-based SmoIP assay allows direct detection of Smo phosphorylation* To develop a system for the direct investigation of Smo activation in cell culture, we chose the *Drosophila* S2 cell line ^29^. These cells express all relevant upstream components of the Hh pathway, but lack the downstream transcription factor Ci and are therefore unable to affect Smo activation through transcriptional feedback ^30^. Epression of the SmoIP sensor from the endogenous promoter yielded insufficient fluorescent signal. To obtain the expression levels required for reliable detection we turned to reporter overexpression, re-establishing a corresponding polyclonal line expressing SmoIP under control of a UAS promoter driven by a tubulin-Gal4VP16 cassette on the same plasmid. We also switched to hygromycin selection, which was expected to produce a higher, average transgene copy number per cell ^26^. Indeed, stimulation of S2 cells transfected with UAS-smoIP tub-Gal4VP16 (in the following abbreviated as UAS-smoIP) with Hh conditioned medium produced a fluorescent signal readily detectable by FACS. However, the proportion of transgenic cells in the population was too low for robust analyses (Fig. 1b). To increase the fraction of transgenic cells in the population we iteratively sorted and recultured the 10% of cells with highest response to Hh stimulation based on sensor fluorescence. Already after two rounds of sorting about 90% of cells were responsive (Fig 1c). Since P-element based transgenesis produces random integration in the genome we decided to subclone individual cells to produce genetically homogeneous lines with the desired response properties for use in subsequent experiments. Using a soft agar cloning technique in combination with an irradiated feeder cell layer ^27^ we obtained and analyzed 20 individual clones harboring UAS-SmoIP insertions. For the following analysis we selected clone 14 (cl14), which showed the highest ratio of induction signal over background pathway activity (Fig. 1d, e). As a result of these preparatory experiments we had therefore obtained a stable, genetically homogeneous clone of UAS-SmoIP cells that reproducibly responded to Hh stimulation with a robustly detectable increase in sensor fluorescence.

*UAS-SmoIP fluorescence responds as predicted to experimental manipulation*

To validate that the cl14 assay correctly reflects endogenous Hh pathway behavior we tested system response to perturbation of known components of the Hh signalling cascade. PKA is the main kinase that phosphorylates Smo and promotes its activation. Consistently, the small molecule PKA inhibitor h89 ^31^ completely abolished SmoIP fluorescence in response to Hh treatment (Fig. 2a, b). To test whether PKA activity was sufficient for the full activation of our assay we treated cells with two PKA activators, forskolin (fsk) and IBMX, that had both previously been shown to work in *Drosophila* S2 cells ^32, 33^. Even though both inhibitors activate PKA by increasing intracellular cAMP levels, they do so by different mechanisms: While fsk stimulates adenylate cyclase activity, IBMX inactivates phosphodiesterases. Importantly, the two PKA activators exhibited consistent effects on SmoIP sensor fluorescence: In both cases, application of the drug had barely any effect on mock treated cl14 cells, but showed pronounced, cooperative activation when acting together with Hh (Fig. 2a, c, d), reproducing previously published result for fsk ^32^. In contrast, global inactivation of phosphatases by okadaic acid (OA) is sufficient to drive Hh signalling in the basence of ligand ^34^. Consistently, OA treatment by itself induced a sensor response exceeding the fluorescence induced by stimulation with Hh alone (Fig. 2a,e). Interestingly, combined treatment of cl14 cells with OA and Hh did not increase UAS-smoIP response strength over OA alone. In contrast, treatment with fsk or IBMX in combination with OA had a pronounced cooperative effect on Smo phosphorylation that could not be further increased by addition of Hh. These observations suggest that, in the absence of Hh, phosphatases sensitive to OA continuously act to maintain Smo in an inactive state, while the same Smo pool is not accessible by the endogenous PKA, even when the kinase is experimentally activated. Instead, PKA appears to require some prior action on Smo, provided endogenously by Hh or experimentally by OA, before it can promote Smo phosphorylation (Fig. 2a, f, g).

**Figure 2.**
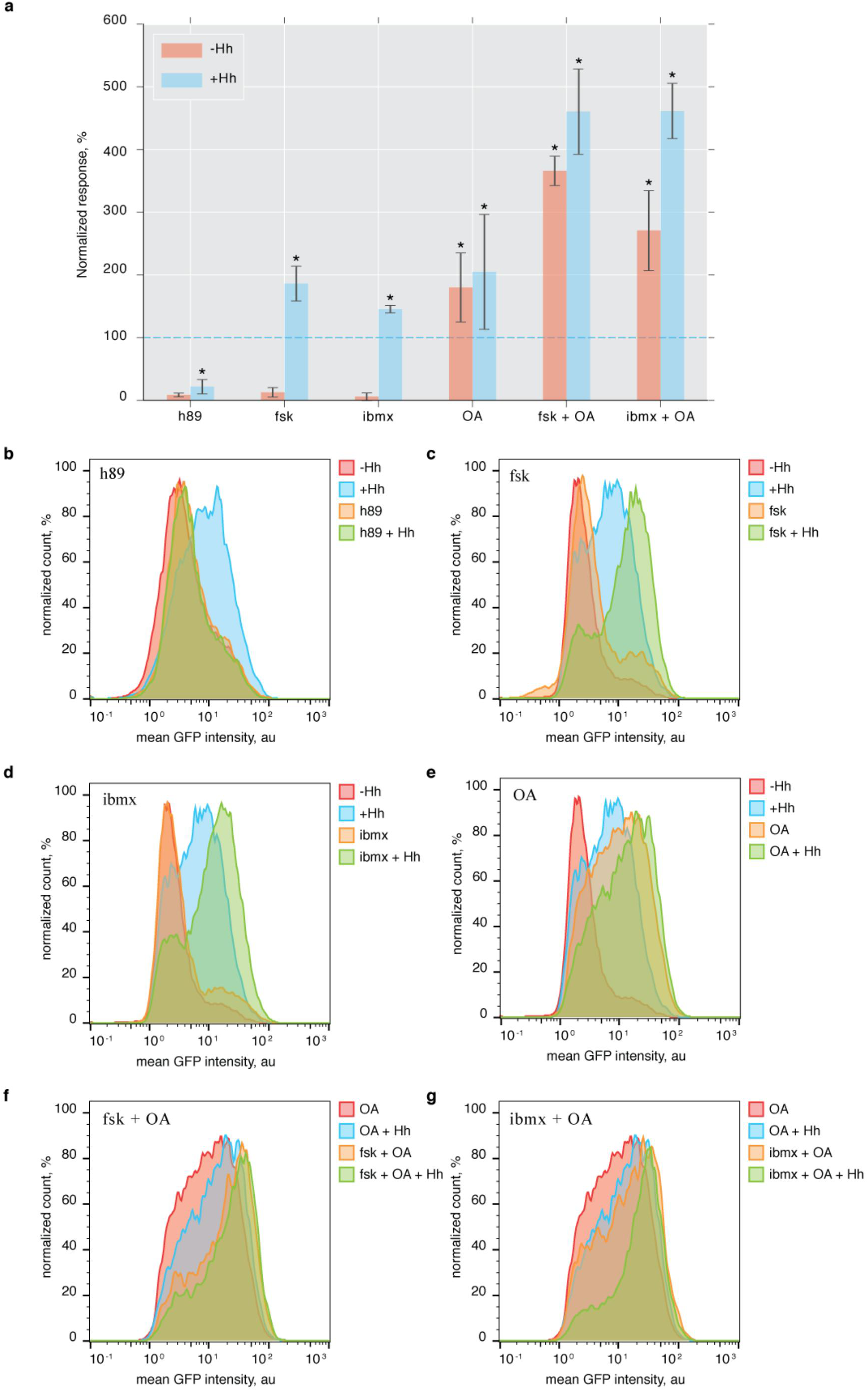
The SmoIP assay captures the endogenous pathway response to chemical perturbation. **(A)** Average effect of h89 (PKA inhibitor), fsk, IBMX (PKA activators), and OA (phosphatase inhibitor), alone or in combination, on baseline and Hh induced SmoIP fluorescence response. Effect of Hh stimulation on untreated cells set as 100% (dashed line). N = 3−4 replicates per experiment, mean ± sd, *p<0.05, m-w test, significant change compare to control Hh stimulation. (B-G) FACS histograms for individual experiments using the indicated components.

As a final test of assay specificity we knocked down several pathway regulators by transfection with dsRNA (Supplementary Fig. S1). Knockdown of Smo, which also eliminates the sensor itself, and of Fused (Fu), a Hh pathway member acting downstream of Smo, were used as positive and negative controls, respectively. As expected, transfection of Fu dsRNA had no effect, while Smo dsRNA reduced sensor fluorescence. Conversely, knockdown of Ptc mimicked stimulation with Hh, leading to sensor activation even in the absence of ligand. Protein stabilization is known to play a major role in Hh signal transduction. Degradation of Smo via the proteasome/lysosome pathway ensures that signalling is kept down in the absence of ligand ^13,14^. Accordingly, knockdown of Uba1, the sole E1 enzyme in *Drosophila*, slightly increased sensor fluorescence in the absence of Hh. Finally, inhibition of endocytosis by knockdown of the *Drosophila* Dynamin homologue Shibire, which we had previously shown to be sufficient to induce Smo phosphorylation ^16^ strongly increased the sensor signal, exceeding the values achieved by stimulation with Hh alone. Therefore, as in the case of chemical perturbation, knockdown of the endogenous signal transduction machinery produced the expected responses in our assay.

Summarizing these experiments we could therefore conclude that the cl14 system correctly reports Smo activation in cell culture.

*Stimulation of cl14 cells with increased Hh levels reveals cell specific response thresholds*

The assay system we had developed and validated gave us an opportunity to determine the Smo response to Hh stimulation quantitatively and with single cell resolution. This quantitative approach promised to shed light on the how target cells may respond to graded Hh signals, a recurring and important theme in developmental biology. We therefore first decided to explore how the cl14 sensor system responded over varying Hh concentrations and stimulation times.

For stimulation of cultured cells a 1:1 mixture of fresh and Hh conditioned medium is typically assumed to provide full pathway activation ^35^. We therefore use this mixing ratio to build the time response curve of our assay. Upon stimulation, the fluorescent sensor signal becomes first detectable after 4h and then gradually increases up to 24h under continuous stimulation (Fig. 3a). This kinetics does not reflect the direct measurement of Smo accumulation rate after Hh stimulation, where an increase of protein level is detectable as soon as 30min after onset of stimulation ^36^. Presumably, the slower kinetics of our system reflects a lower sensitivity of fluorescence measurement compared with immunoblotting, therefore requiring prolonged signal integration and protein stabilization. To maximise the detection window of our assay we fixed stimulation time at 24h throughout all further experiments.

**Figure 3.**
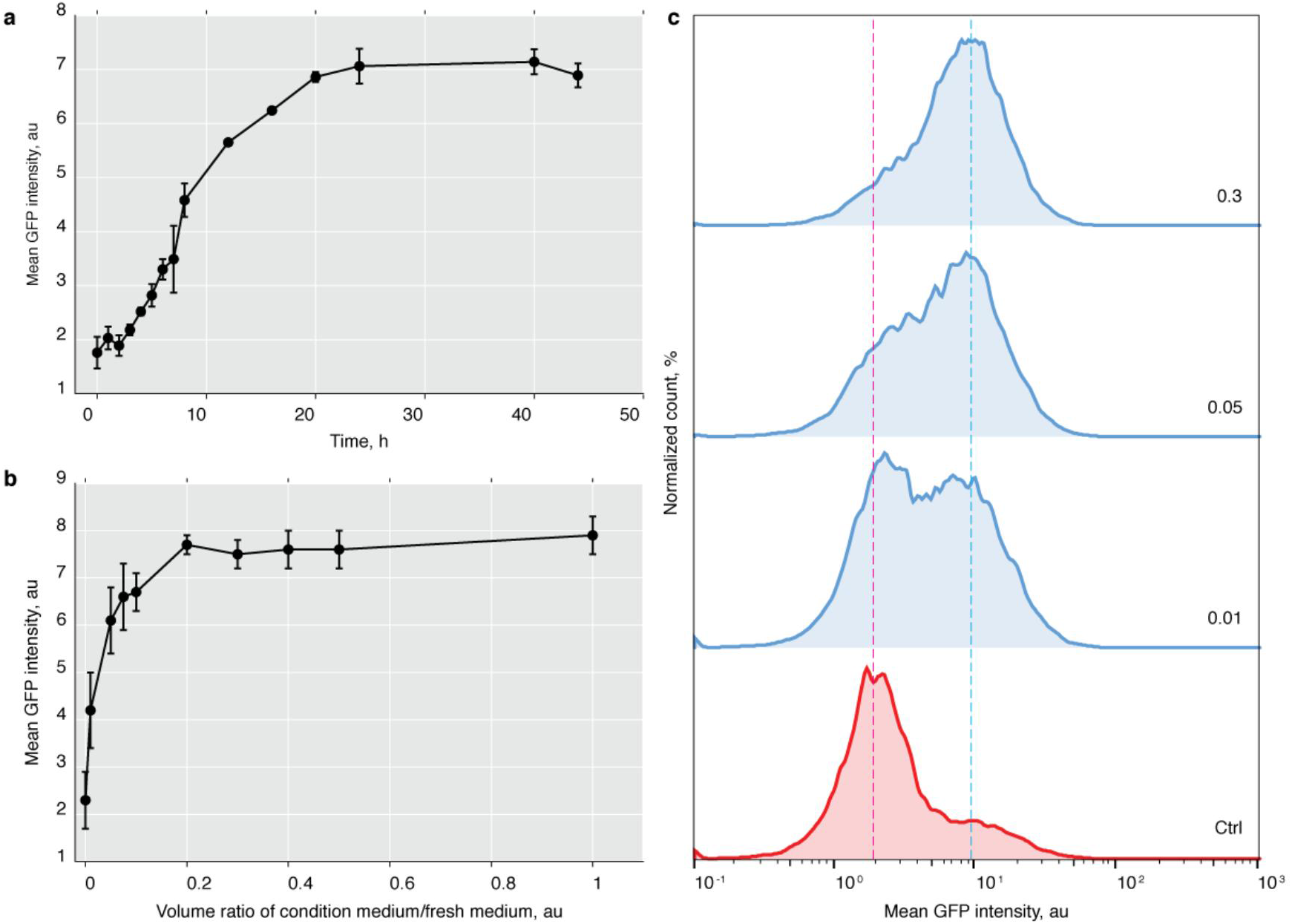
Time and concentration dependence of the SmoIP response. (A) SmoIP fluorescence signal of cl14 cells stimulated 1:1 with Hh conditioned medium plotted against stimulation time (B) SmoIP fluorescence signal of cl14 cells after 24h of stimulation plotted against ratio of Hh conditioned and fresh medium. (C) FACS histograms for individual points of the concentration curve, numbers represent ration between condition and fresh medium. (A, B) N = 2−6 replicates, mean ± sd.

We next explored the volume ratio between conditioned and fresh medium in a range from 1:100 to 1:1 (Fig. 3b). The system reached saturation at ratio around 1:5, however, we were able to detect pathway activation even at the lowest concentration we tested. Taking advantage of the fact that the FACS based assay provides information about signal intensity for each individual cell we broke the average values presented in (Fig. 3b) down to response histograms for several fractions of Hh conditioned medium ranging from 0 to 0.3. Interestingly, at intermediate stimulation levels the average fluorescence signals exhibited a bimodal distribution, comprising of two populations of cells with high and low reporter fluorescence. Rather than shifting both populations towards higher fluorescence levels, increasing Hh levels caused a larger fraction of the cells to phosphorylate the SmoIP sensor (Fig. 3c). Even though our cl14 assay line is clonally derived from a single cell, each cell in the tested population thus appears to have a different concentration threshold above which it activates its Hh signalling cascade.

*A small molecule inhibitor screen suggests an effect of RTK signalling and cell cycle associated kinases on Smo activation*

The real power of *in-vitro* assays such as the one we developed lies in the possibility to perform high throughput (HT) experiments. We therefore decided to optimize our assay for future HT studies. This was facilitated by the fact that S2 cells can be cultivated in suspension, which allowed us to easily implement FACS as a detection system and thus immediately obtain quantitative data. To minimize screening time, a typical problem of FACS based assays, and to increase throughput, we scaled the assay down to 384 well plate format.

As proof of principle we systematically tested a Selleckchem collection of 1130 small molecule inhibitors with known molecular targets (Selleckchem L1100) for effects on Smo phosphorylation state. However, these inhibitors were developed and validated for mammalian systems, and for the majority of them there is no information about their efficacy in *Drosophila*. We therefore screened each compound at two concentrations, 2*µ*m and 15*µ*M. Cells were pre-incubated with compounds for 30 min, stimulated with either Hh or mock conditioned medium, and analyzed by FACS after 24h (Fig. 4a). Measuring the effects of inhibitors on stimulated and unstimulated fluorescence baselines enabled us in principle to detect both activators and inhibitors of Smo phosphorylation. The effect of each given treatment was normalized to the increase in sensor fluorescence caused by Hh stimulation of otherwise untreated cells, which was set as 100%.

**Figure 4.**
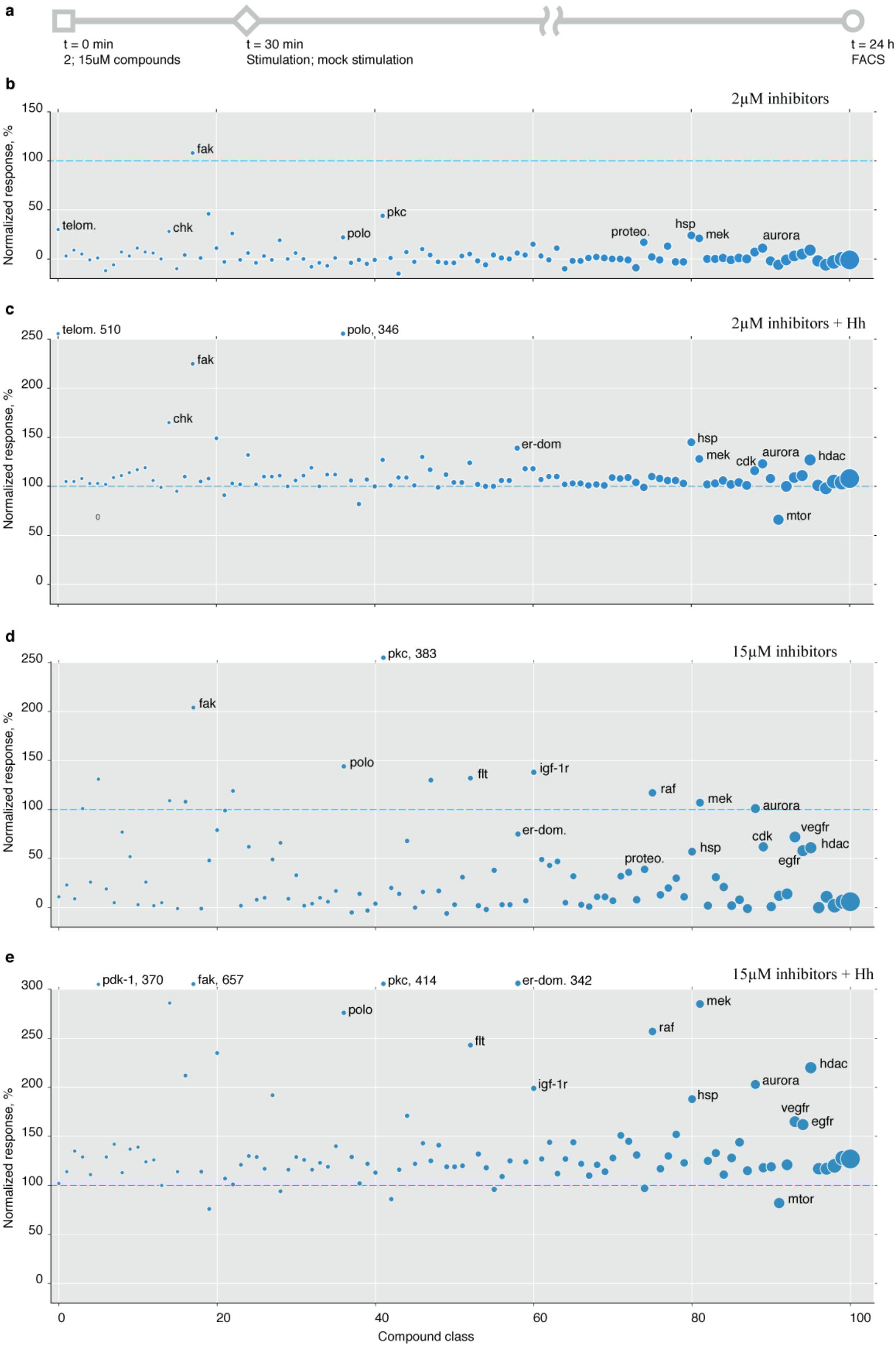
Screening a small molecule inhibitor library for modulation of SmoIP fluorescence. (A) Schematic representation of screen setup. (B-E) Normalized effect of inhibitors grouped according to their primary targets on SmoIP fluorescence. Effect of Hh stimulation on untreated cells set as 100% (dashed line). Diameter of circle reflects number of components per cluster, cutoff N ≥ 3, clusters sorted along X axis accordingly. (B, C), inhibitors used at 2*µ*M, (D, E) at 15*µ*M.

Surprisingly, we found few compounds capable of downregulating Smo phosphorylation upon stimulation with Hh. In contrast, many more compounds increased Smo phosphorylation, producing a continuous spectrum of responses ranging from mild increases up to 10-20 times over control (Supplementary Fig. S2, Supplementary table S1). We therefore clustered the compounds according to their molecular targets and computed median response for each of these classes (Fig. 4b-e, Supplementary table S2).

Hits were then identified as clusters where multiple inhibitors targeting one or more components of the same pathway exhibited a consistent effect on sensor fluorescence. Correlating the observed effect for each such cluster between experiments performed at different concentrations and in the presence vs. absence of Hh stimulation demonstrated reproducibility and robustness of our assay (Supplementary Fig. S2).

Clustering compounds by targets in this way revealed several candidate pathways whose inhibition appeared to modulate Smo activation. Sensor fluorescence was increased by compounds targeting various RTKs (Fig. 4b-e) or the downstream Raf and MAPK signalling complexes (Figs. 4b-e and 5). A second cluster exhibiting an increased SmoIP signal was comprised of compounds targeting kinases regulating cell cycle progression and microtubule organization (polo, polo-like, and aurora kinases) (Fig. 4b-e and 5). A third cluster of compounds increasing sensor fluorescence consisted of inhibitors of histone deacetylases (Fig. 4b-e and 5). The sole class of inhibitors consistently downregulating sensor response to Hh was the group of mTOR inhibitors: More than half of these compounds (15/23) reduced the fluorescence response to Hh stimulation by at least 20% (Fig. 4b-e and 5).

**Figure 5.**
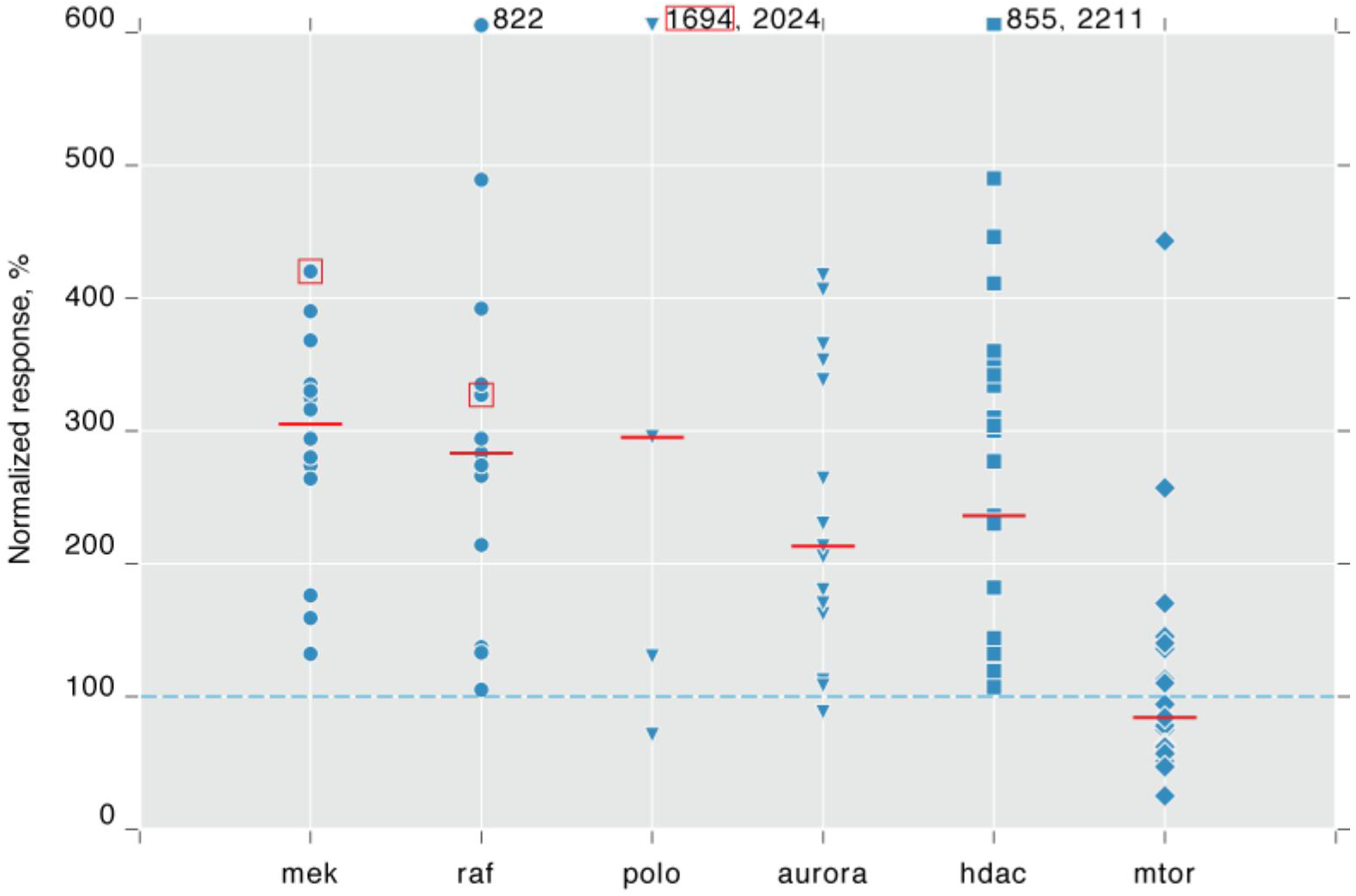
Inhibitor classes with consistent effect on Smo activation. SmoIP fluorescence of cl14 cells stimulated for 24h with Hh conditioned medium in the presence of individual compounds inhibiting the indicated, selected primary targets. Effect of Hh stimulation on untreated cells set as 100% (dashed line). Inhibitor concentration 15*µ*M, red bars indicate median, boxes mark compounds selected for subsequent validation experiments.

Thus, our screen implicated three major signalling pathways and the epigenetic machinery in the regulation of Smo phosphorylation, neither of which had previously been shown to act near the top of the Hh pathway.

*Validation of hits via independent, secondary assays confirms the links between MAPK pathway or polo kinase inhibitors and Smo activation.*

The mTOR protein is part of MTORC1 complex that acts as a central regulator of cell growth and protein synthesis ^37^. Since Smo protein is constantly turned over, its accumulation in response to Hh, especially over the long stimulation period of our assay, involves *de novo* protein synthesis. The negative role of mTOR inhibition on SmoIP sensor accumulation was therefore not unexpected, and we decided to focus instead on the compound clusters causing increased sensor fluorescence. We also decided to exclude the cluster of HDAC inhibitors, as epigenetic modifiers most likely affect Smo activation only indirectly.

From the remaining clusters, we selected three highly scoring compounds: To interfere with signalling downstream of RTKs at two different levels we chose the Raf inhibitor dabrafenib (dbn) ^38^ and the MEK inhibitor tak-733 ^39^. Since the two kinases act sequentially within the same signalling cascade, a shared mode of Smo activation by their respective inhibitors would lend credibility to our assay. Volasertib (vlt) ^40^ was used as an example of a polo inhibitor strongly affecting Smo activation (Fig. 5).

We first confirmed that incubation of cl14 cells with either compound for 24h did not lead to increased cell death or abnormal cell morphologies observed by changes in FACS forward and side scatter or by microscopy. The selected compounds thus do not appear to cause nonspecific cytotoxicity. Second, we repeated the FACS experiments at larger scale and tested the induction of sensor fluorescence by live cell microscopy. All three compounds stimulated fluorescence of the SmoIP sensor at levels higher than Hh treatment alone (Fig. 6a) and caused accumulation of activated fluorescent SmoIP sensor at the cell periphery (Fig. 6b). Induction of sensor fluorescence by all compounds was concentration dependent (Supplementary Fig. S3), further validating the initial observations made in HT mode and suggesting that the observed effects were specific.

**Figure 6.**
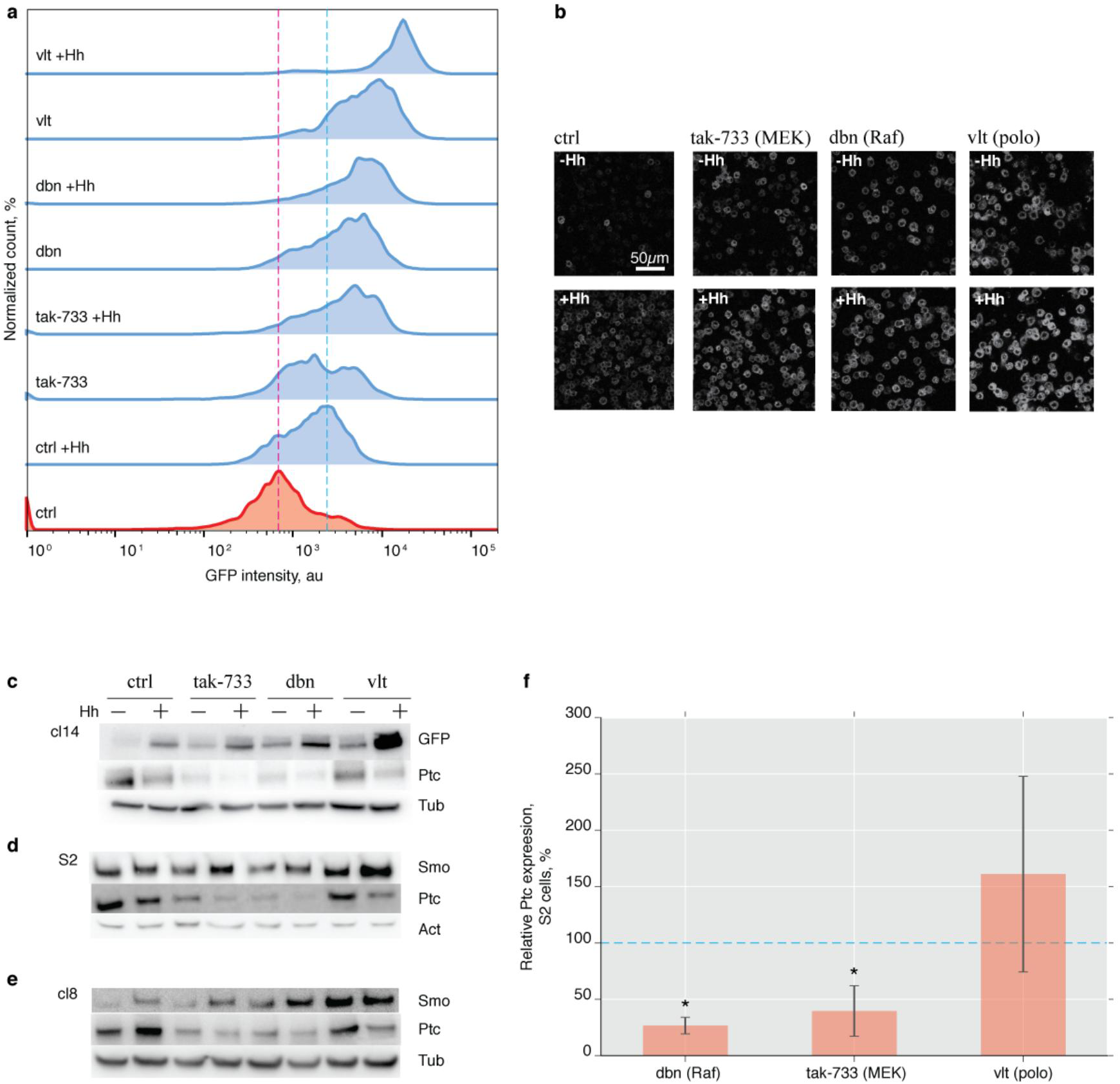
Validation of screen results for selected inhibitors. (A) FACS histogramsof SmoIP fluorescence following treatment with 15*µ*M tak-733 (MEK inhibitor), dobrafenib (dbn, Raf inhibitor) or volasertib (vlt, polo inhibitor) in the presence or absence of Hh. Red and blue dashed line indicate baseline and Hh induced fluorescence, respectively, in untreated control cells. (B) Confocal live-cell images corresponding to the histograms in (A). (C-E) Western blots for GFP, Smo and Ptc on lysates of Cl14 (C), S2 (D) and cl8 cells (E) in the presence of indicated inhibitors. Tubulin (C, E) or actin (D) used as loading control. (F) Ptc transcription in S2 cells following inhibitor treatment. Expression normalized to GAPDH2, N = 3−4, mean ± sd, *p<0.05, m-w test. (B) Scale bar, 50 *µ*m.

However, assay conditions, including expression levels and stimulation times, were optimized to achieve the maximal fluorescence signal. Phosphorylation of the sensor does therefore not necessarily directly reflect activation of the endogenous signal cascade. We therefore validated our FACS and microscopy results by methodologically independent biochemical experiments in our cl14 reporter cell line (Fig. 6c), as well as in wildtype S2 cells (Fig. 6d) and in cl8 cells (Fig. 6e), an unrelated cell line that expresses the entire Hh signalling cascade including the transcription factor Ci and is thus endogenously sensitive to Hh ^30^. Stimulation with hh conditioned medium or inhibitor treatment of cl14 cells individually led to increased SmoIP protein levels. This response was further enhanced by combining ligand and inhibitors (Fig. 6c). A similar effect, albeit with a reduced dynamic range, was also seen for endogenous Smo protein in S2 cells (Fig. 6d) and in cl8 cells (Fig. 6e). Thus, activation and accumulation of Smo in response to these compounds is not merely an artefact of our fluorescent assay system. In addition, extracellular labelling of cl14 reporter cells using an antibody recognizing the Smo N-terminus confirmed Smo translocation to the plasma membrane in response to stimulation with Hh or compound treatment, another hallmark of Hh pathway activation (Supplementary Fig. S4).

Thus, all three inhibitors selected for the validation experiments activate the Hh signalling cascade, at least down to the level of Smo phosphorylation and accumulation at the plasma membrane.

*MAPK pathway and polo inhibitors activate Hh signalling by distinct mechanisms*

The successful validation of these hits encouraged us to investigate the underlying mechanisms of Smo activation. The Hh receptor Ptc is the main, endogenous inhibitor of Smo activation and hence a prime candidate for mediating the observed effect. In cl14 cells and the S2 cells they were derived from, Ptc levels dropped in response to Hh treatment (Fig. 6c-d), presumably reflecting the Hh induced internalization and degradation. However, Ptc is also a target gene of the Hh pathway. Hence, Hh stimulation caused an increase in Ptc levels in the endogenously signalling competent cl8 cells (Fig. 6e). Ptc levels decreased in all three tested cell types following treatment with tak-733 or dbn (Fig. 6c-e). We excluded that these inhibitors inadvertently induced Hh expression (Supplementary Fig. S5). Instead, the reduction in Ptc protein levels in response to dbn and tak-733 was correlated with a reduction in Ptc transcription (Fig. 6f), which may suffice to explain the greater baseline activation and increased Hh sensitivity of the cells treated with the two inhibitors. Smo would in this case still be activated by the endogenous signalling machinery, resembling e.g. pathway activation by Ptc RNAi knockdown.

In contrast, treatment with the polo inhibitor vlt did not affect basal Ptc protein levels in either of the three cell lines (Fig. 6c-e) nor Ptc transcription in S2 cells (Fig. 6f). At the protein level, treatment of unstimulated cl14 cells with vlt stabilized SmoIP comparable to treatment with Hh, even though the induced reporter fluorescene was substantially stronger. Co-stimulation with vlt and Hh further increased both reporter fluorescence and accumulation of SmoIP (Fig. 6c). The same effect was seen for endogenous Smo in S2 cells (Fig. 6d). In the signaling competent cl8 cells, vlt also induced a strong increase in basal Smo levels. However, the pronounced cooperativity with Hh was absent (Fig. 6e). Treating cl8 cells with Hh upregulated Ptc levels, confirming signal transduction down to the transcription level. Importantly, this was not seen following treatment with vlt, despite the observed stabilization of Smo (Fig. 6e). Thus vlt, unlike dbn or tak-733, does not act through activation of the endogenous signalling machinery, but influences Smo phosphorylation and accumulation through a mechanism that does not activate downstream signalling. Consistent with the notion that dbn/tak-733 and vlt act through different mechanisms, the PKA activators IBMX and fsk, which are able to enhance fluorescence of endogenously activated Smo (Fig. 2, Supplementary Fig. S5), show no such cooperative effect with vlt (Supplementary Fig. S5). In contrast, co-treatment with the phosphatase inhibitor OA enhances Smo activation by vlt both at the protein and fluorescence level (Supplementary Fig. S5).

In summary, the small molecule inhibitors dbn and tak-73 that target Raf and MEK, respectively, appear to downregulate Ptc expression in a Ci independent manner, thereby activating Smo through the endogenous signalling machinery. In contrast, the polo inhibitor vlt promotes Smo phosphorylation and accumulation via a different mechanism that fails to trigger downstream signal transduction. While our present experiments cannot dissect these mechanisms in detail, this clearly demonstrates that the assay is in principle able to uncover Smo activation independent of the underlying molecular mechanism, and is thus fit for purpose.

## Discussion

We have presented a novel, FACS based HTS system in *Drosophila* S2 cells that relies on visualizing Smo phosphorylation with the help of a fluorescent sensor. Using our system, we could follow the Smo response to different Hh concentrations quantitatively and on the single cell level. Furthermore, by screening a library of small molecule inhibitors, we identified groups of compounds with shared molecular targets that affected Smo phosphorylation. Our approach thus provides a valuable tool for the systematic and quantitative investigation of upstream events in the Hh signaling pathway.

Monitoring the degree of Smo activation for every cell in a large population provides a quantitative resolution that cannot be obtained by bulk, biochemical methods, transcriptional reporters, or imaging based approaches restricted to smaller sample sizes. Pathway response to increasing ligand concentrations is a key problem in developmental biology, where Hh morphogen gradients are interpreted to drive differential target gene expression. According to one model, increasing Hh stimulation increases the number phosphorylated residues in each Smo molecule ^22^, presumably by sequential recruitment of distinct serine clusters ^41^. Bulk assays such as immunoblots with phospho-specific Smo antibodies indeed suggested such a graded Smo phosphorylation response ^41^. However, our experiments revealed that this quantitative response is apparently not due to gradually increasing Smo phosphorylation levels in each cell. Instead, submaximal Hh level only induced Smo phosphorylation in a subset of all potential target cells. Higher Hh levels increased the ratio of responding to non-responding cells, but did not increase absolute fluorescence levels in the responders. Thus, even in our clonally derived population, individual cells must have different thresholds for Smo activation. Importantly, all cells in the population were, in principle, capable of responding to the Hh upon stimulation at saturating Hh concentrations. It will be interesting to see whether this variation in sensitivity, both between cells or potentially even within one cell over time, also occurs in tissues *in vivo*.

A second, interesting observation concerns the activation of Smo by PKA. Depending on the presence or absence of Hedgehog, Smo forms biochemically detectable complexes with PKA ^32^ or phosphatases ^34^. Expression of constitutively active murine PKA *in vivo* is by itself sufficient to induce fluorescence of the SmoIP reporter and Hh target gene expression ^16^, but activation of the endogenous PKA pool by pharmacologically increasing cAMP levels through fsk or IBMX treatment is, at least *in vitro*, incapable of activating Smo. However, both components strongly increase Smo activation in the presence of Hh, suggesting that the activated, endogenous kinases can under these condition act on the Smo sensor. In contrast, the phosphatase inhibitor OA does not require cooperative action of Hh, suggesting that its target phosphatases PP1 and PP2A can access Smo in unstimulated cells. These observations are readily compatible with a constraint we identifed during mathematical simulation of Smo activation ^16^, whereby PKA activity must be restricted to the plasma membrane, where it acts on Smo trafficked there following addition of ligand.

In addition to providing quantiative data from single experiments, our assay is scaleable to high throughput mode, up to 384 well plates. We performed a proof of principle screen using a library of small molecules with known molecular targets. Grouping compounds by these targets revealed several compound clusters whose members consistently affected Smo phosphorylation. The emergence of these clusters by itself provides a strong validation for the screening results, even though we are not able to extrapolate from the available information on the individual compounds to specific targets in the fly system. Although it was beyond the scope of this study to identify the primary targets of these hits we were able to define a putative mechanism of action for two of the selected chemicals: Inhibition of the MAPK cascade by dbn (Raf) and tak-733 (MEK) downregulated Ptc transcription, thus presumably promoting Smo activation by derepression of the endogenous signalling machinery. This provides a novel tool to manipulate Ci-independent Ptc expression. In contrast, treatment of our reporter cells with vlt, a presumptive polo inhibitor, induced Smo phosphorylation, accumulation, and relocalisation to the plasma membrane independent of Ptc inhibition. However, this was not associated with signal transduction downstream of Smo, even in the fully Hh sensitive cl8 cells. Importantly, neither mode of Smo activation could have been uncovered by a screen using a transcriptional readout, which in the case of the Hh pathway are usually based on the Ptc promotor, highlighting the benefits of using multiple, complementary approaches.

Taken together, our experimental results validated the SmoIP assay as a powerful tool for investigating Smo activation that can be employed in high-throughput analysis and yields quantitative data for the investigation of individual cell responses.

## Acknowledgements

We thank Michal Zurovec, the DGRC, Steve Brown and the Sheffield RNAi screening facility, and Suzanne Eaton for advice and reagents. We thank the CRTD FACS facility and the MPI-CBG screening facility for their support, and Jared Sterneckert for providing the library, discussions and critical review of the manuscript. The project was supported by a CRTD seed grant and DFG grant BO 3270/4-1 to CB.

### Author contributions

EAA designed, performed and analyzed the experiments and wrote the manuscript, CB initiated the project, analyzed the data, and wrote the manuscript.

### Competing financial interests

Authors declare no competing financial interests

## References

1 Briscoe, J. & Therond, P. P. The mechanisms of Hedgehog signalling and its roles in development and disease. Nat Rev Mol Cell Biol 14, 416–429, (2013).

2 Kuwabara, P. E. & Labouesse, M. The sterol-sensing domain: multiple families, a unique role? Trends in genetics : TIG 18, 193–201, (2002).

3 Corcoran, R. B. & Scott, M. P. Oxysterols stimulate Sonic hedgehog signal transduction and proliferation of medulloblastoma cells. Proc Natl Acad Sci U S A 103, 8408–8413, (2006).

4 Taipale, J., Cooper, M. K., Maiti, T. & Beachy, P. A. Patched acts catalytically to suppress the activity of Smoothened. Nature 418, 892–897, (2002).

5 Nachtergaele, S. et al.Oxysterols are allosteric activators of the oncoprotein Smoothened. Nat Chem Biol 8, 211–220, (2012).

6 Byrne, E. F. et al.Structural basis of Smoothened regulation by its extracellular domains. Nature 535, 517–522, (2016).

7 Huang, P. et al.Cellular Cholesterol Directly Activates Smoothened in Hedgehog Signaling. Cell 166, 1176–1187e1114, (2016).

8 Khaliullina, H., Bilgin, M., Sampaio, J. L., Shevchenko, A. & Eaton, S. Endocannabinoids are conserved inhibitors of the Hedgehog pathway. Proc Natl Acad Sci U S A 112, 3415–3420, (2015).

9 Yavari, A. et al.Role of lipid metabolism in smoothened derepression in hedgehog signaling. Dev Cell 19, 54–65, (2010).

10 Jiang, K. et al.PI(4)P Promotes Phosphorylation and Conformational Change of Smoothened through Interaction with Its C-terminal Tail. PLoS Biol 14, e1002375, (2016).

11 Jia, J., Tong, C., Wang, B., Luo, L. & Jiang, J. Hedgehog signalling activity of Smoothened requires phosphorylation by protein kinase A and casein kinase I. Nature 432, 1045–1050, (2004).

12 Li, S. et al.Regulation of Smoothened Phosphorylation and High-Level Hedgehog Signaling Activity by a Plasma Membrane Associated Kinase. PLoS Biol 14, e1002481, (2016).

13 Li, S. et al.Hedgehog-regulated ubiquitination controls smoothened trafficking and cell surface expression in Drosophila. PLoS Biol 10, e1001239, (2012).

14 Xia, R., Jia, H., Fan, J., Liu, Y. & Jia, J. USP8 promotes smoothened signaling by preventing its ubiquitination and changing its subcellular localization. PLoS Biol 10, e1001238, (2012).

14 Ma, G. et al.Regulation of Smoothened Trafficking and Hedgehog Signaling by the SUMO Pathway. Dev Cell 39, 438–451, (2016).

16 Kupinski, A. P. et al.Phosphorylation of the Smo tail is controlled by membrane localization and is dispensable for clustering. J Cell Sci, (2013).

17 Lum, L. et al.Identification of Hedgehog pathway components by RNAi in Drosophila cultured cells. Science 299, 2039–2045, (2003).

18 Nybakken, K., Vokes, S. A., Lin, T. Y., McMahon, A. P. & Perrimon, N. A genome-wide RNA interference screen in Drosophila melanogaster cells for new components of the Hh signaling pathway. Nat Genet 37, 1323–1332, (2005).

19 Frank-Kamenetsky, M. et al.Small-molecule modulators of Hedgehog signaling: identification and characterization of Smoothened agonists and antagonists. Journal of biology 1, 10, (2002).

20 Chen, J. K., Taipale, J., Young, K. E., Maiti, T. & Beachy, P. A. Small molecule modulation of Smoothened activity. Proc Natl Acad Sci U S A 99, 14071–14076, (2002).

21 Schaefer, G. I. et al.Discovery of small-molecule modulators of the Sonic Hedgehog pathway. Journal of the American Chemical Society 135, 9675–9680, (2013).

22 Zhao, Y., Tong, C. & Jiang, J. Hedgehog regulates smoothened activity by inducing a conformational switch. Nature 450, 252–258, (2007).

23 Nagai, T., Sawano, A., Park, E. S. & Miyawaki, A. Circularly permuted green fluorescent proteins engineered to sense Ca2+. Proc Natl Acad Sci U S A 98, 3197–3202, (2001).

24 Jordan, M. & Wurm, F. Transfection of adherent and suspended cells by calcium phosphate. Methods 33, 136–143, (2004).

25 Chou, T. B., Noll, E. & Perrimon, N. Autosomal P[ovoD1] dominant female-sterile insertions in Drosophila and their use in generating germ-line chimeras. Development 119, 1359–1369, (1993).

26 Segal, D., Cherbas, L. & Cherbas, P. Genetic transformation of Drosophila cells in culture by P element-mediated transposition. Somat Cell Mol Genet 22, 159–165, (1996).

27 Echalier, G. Drosophila Cells in Culture. (Academic Press, 1997).

28 Issigonis, M. & Matunis, E. The Drosophila BCL6 homolog Ken and Barbie promotes somatic stem cell self-renewal in the testis niche. Developmental biology 368, 181–192, (2012).

29 Schneider, I. Cell lines derived from late embryonic stages of Drosophila melanogaster. Journal of embryology and experimental morphology 27, 353–365, (1972).

30 Cherbas, L. et al.The transcriptional diversity of 25 Drosophila cell lines. Genome Res 21, 301–314, (2011).

31 Chijiwa, T. et al.Inhibition of forskolin-induced neurite outgrowth and protein phosphorylation by a newly synthesized selective inhibitor of cyclic AMP-dependent protein kinase, N-[2-(p-bromocinnamylamino)ethyl]-5-isoquinolinesulfonamide (H-89), of PC12D pheochromocytoma cells. The Journal of biological chemistry 265, 5267–5272, (1990).

32 Li, S., Ma, G., Wang, B. & Jiang, J. Hedgehog induces formation of PKA-Smoothened complexes to promote Smoothened phosphorylation and pathway activation. Science signaling 7, ra62, (2014).

33 Wang, B. et al.The insulin-regulated CREB coactivator TORC promotes stress resistance in Drosophila. Cell metabolism 7, 434–444, (2008).

34 Jia, H., Liu, Y., Yan, W. & Jia, J. PP4 and PP2A regulate Hedgehog signaling by controlling Smo and Ci phosphorylation. Development 136, 307–316, (2009).

35 Zhang, Y. et al.Transduction of the Hedgehog signal through the dimerization of Fused and the nuclear translocation of Cubitus interruptus. Cell research 21, 1436–1451, (2011).

36 Zhu, A. J., Zheng, L., Suyama, K. & Scott, M. P. Altered localization of Drosophila Smoothened protein activates Hedgehog signal transduction. Genes Dev 17, 1240–1252, (2003).

37 Ma, X. M. & Blenis, J. Molecular mechanisms of mTOR-mediated translational control. Nat Rev Mol Cell Biol 10, 307–318, (2009).

38 Menzies, A. M., Long, G. V. & Murali, R. Dabrafenib and its potential for the treatment of metastatic melanoma. Drug design, development and therapy 6, 391–405, (2012).

39 Dong, Q. et al.Discovery of TAK-733, a potent and selective MEK allosteric site inhibitor for the treatment of cancer. Bioorganic & medicinal chemistry letters 21, 1315–1319, (2011).

40 Rudolph, D. et al.BI 6727, a Polo-like kinase inhibitor with improved pharmacokinetic profile and broad antitumor activity. Clinical cancer research : an official journal of the American Association for Cancer Research 15, 3094–3102, (2009).

41 Fan, J., Liu, Y. & Jia, J. Hh-induced Smoothened conformational switch is mediated by differential phosphorylation at its C-terminal tail in a dose-and position-dependent manner. Developmental biology 366, 172–184, (2012).

